# Migratory hoverflies orientate north during spring migration

**DOI:** 10.1101/2022.08.08.503145

**Authors:** Will L. S. Hawkes, Scarlett T. Weston, Holly Cook, Toby Doyle, Richard Massy, Eva Jimenez Guri, Rex E. Wotton Jimenez, Karl R. Wotton

## Abstract

Migratory hoverflies are long-range migrants that, in the northern hemisphere, move seasonally to higher latitudes in the spring and lower latitudes in the autumn. The preferred migratory direction of hoverflies in the autumn has been the subject of radar and flight simulator studies while spring migration has proved to be more difficult to characterise due to a lack of ground observations. Consequently, the preferred migratory direction during spring has only been inferred from entomological radar studies and patterns of local abundance and currently lacks ground confirmation. Here, during a springtime arrival of migratory insects onto the Isles of Scilly and mainland Cornwall, UK, we provide ground proof that spring hoverfly migrants have an innate northward preference. Captured migratory hoverflies displayed northward vanishing bearings when released under sunny conditions under both favourable wind and zero-wind conditions. In addition, and unlike autumn migrants, spring individuals were also able to orientate when the sun was obscured. Analysis of winds suggests an origin for insects arriving on the Isles of Scilly in from western France. These findings of spring migration routes and preferred migration directions are likely to extend to the diverse set of insects found within the western European migratory assemblage.

## 1. Background

Insects migrate in their trillions globally every year to exploit seasonal resources, improve their reproductive success, and/or escape deteriorating habitats (1–4). Insect movements spread vital ecological roles over large areas including pollination, pest control, decomposition, and nutrient transfer (1,4–6). Various approaches have been employed to investigate their orientation during migration including vertical looking radars to monitor insects at high altitudes, flight simulator experiments to assess orientation in a controlled environment and vanishing bearings to assess butterfly orientation within a natural setting (1,7–12). Radar studies in migratory hoverflies have demonstrated seasonally favourable directions of movement, south in autumn and north in spring (7). In addition, during the autumnal migratory season, migratory hoverflies caught at ground level and flown in a flight simulator have been shown to use a time-compensated sun compass to orientate south but fail to orientate when the sun is obscured (11). Equivalent ground evidence is currently lacking for spring hoverfly migrants.

To investigate if spring migratory hoverflies have an innate direction preference, and to see if this direction is influenced by the visibility of the sun or wind direction, we undertook a series of experiments to measure the vanishing bearings of captured, pre-reproductive, migratory hoverflies following their arrival into Cornwall and the Isles of Scilly, UK, in mid-June 2022. We predict that: (1) hoverflies would continue to orientate northwards in a seasonally favourable direction when given a view of the sky with the sun; (2) that they will lose this ability when the sun is obscured; and (3) that they will utilise favourable winds to aid their flight north.

## 2. Methods

### (a) Migratory insect arrival and capture

In 2022 a large influx of migratory insects, consisting of Diptera and Lepidoptera, appeared on the Isles of Scilly and mainland Cornwall between the 16th and 21^st^ June. We captured 66 migratory hoverflies and performed vanishing bearing experiments on the 16^th^ (n=4) and 17^th^ (n=25) in the Isles of Scilly. Following these island experiments we conducted further vanishing bearing experiments in Cornwall on the 20^th^ (n=20) and 21^st^ (n=17). Five migratory species were caught: *Syrphus vitripennis* (n = 45), *Episyrphus balteatus* (n = 13), *Scaeva pyrastri* (n = 4), *Eristalis tenax* (n = 2) and *Syrphus ribesii* (n = 2). Captured hoverflies were placed into a mesh insect cage and experimented upon within 15 minutes of capture.

### (b) Vanishing bearings

Two researchers sat east and west between the cage of captured hoverflies. Hoverflies were removed from the cage by hand, before being held in the air and released above the researcher’s heads. In general, the hoverflies waited >2 seconds before leaving, while on three occasions, the hoverflies left instantly. On departure, both researchers recorded the heading until the hoverfly was no longer visible (approximately 20 meters) and this angle was recorded as the vanishing bearing. Headings were determined using a compass and/or the compass function on a Garmin Instinct Solar watch. Hoverflies were released during three distinct meteorological conditions: (1) Sunny with no wind; (2) sunny with wind (Figure 1a); and (3) during a thick sea mist that obscured the sun and with wind (Figure 1b).

**Figure 1.**
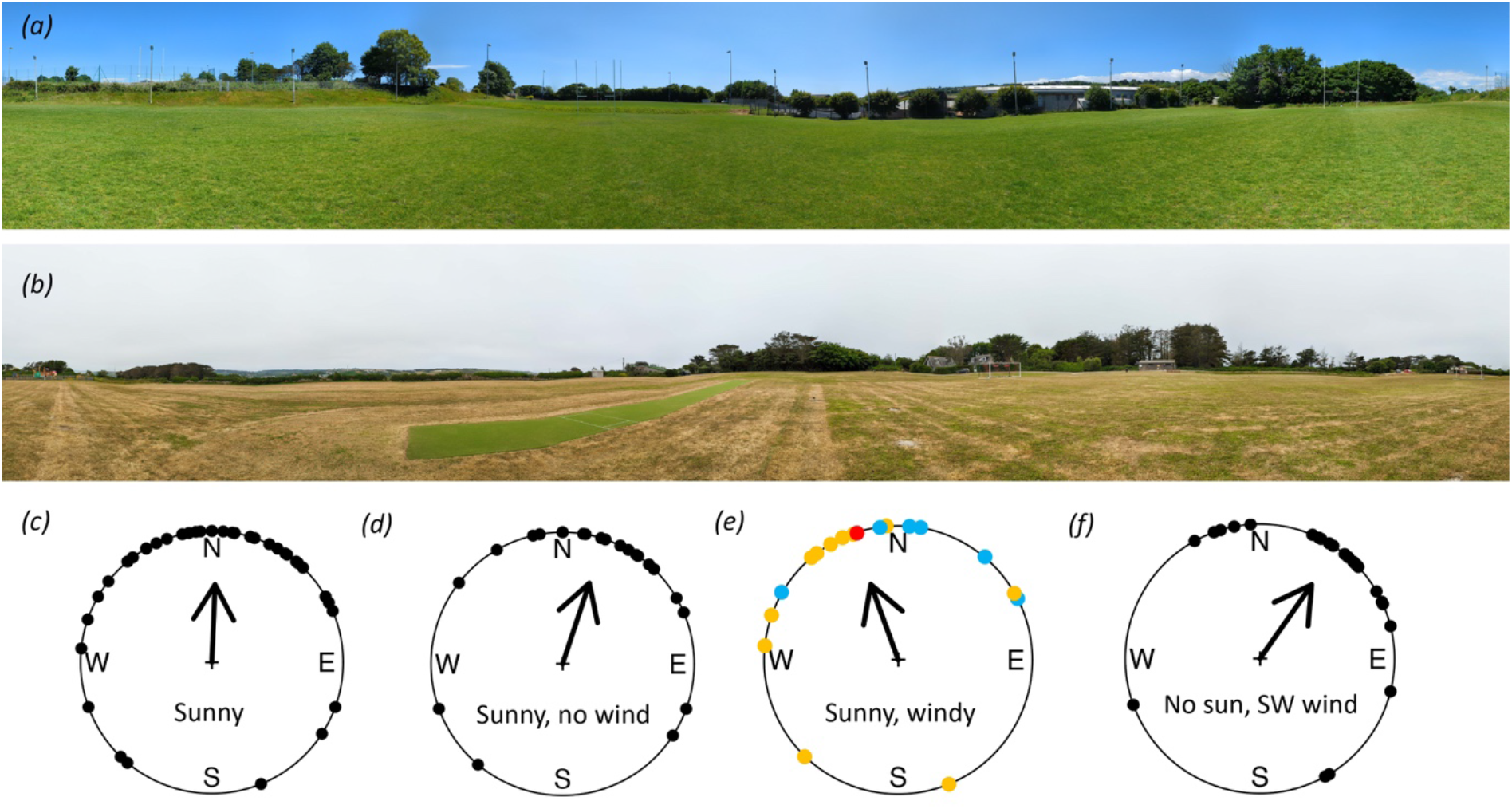
(a) A 360° panoramic photo of the Penryn site on the day of release in sunny conditions. (b) A 360° panoramic photo of the Isles of Scilly study site on the day of release under thick sea mist conditions. (c to f) Circular histograms of individual vanishing bearings (black dots) and group mean directions (black arrow) with length of the arrow depicting *r* from 0 to 1 at the outer edge of the circle. (c) All hoverfly headings while sunny. (d) Hoverfly headings with no wind when sunny. (e) Hoverfly headings while it was windy and sunny. Yellow dots indicate a south-southeasterly wind, blue dots indicate a westerly wind, red dot indicates an easterly wind. (f) Hoverfly headings during a thick sea mist when the sun was obscured with wind from the southwest.

### (c) Location

Experiments took place in two locations: the Isles of Scilly and around the Exeter university campus in Penryn, Cornwall. The Isles of Scilly are an archipelago of islands situated 47km southwest from Land’s End, Cornwall. Experiments were performed on a large open area of a football pitch situated on the Garrison headland (49.913136 N, −6.3211459 W) in the southwest of the island of St Mary’s during a thick sea mist between the hours of 1600 and 1700 (Figure 1b). The study area was a flat expanse 116 m x 177 m surrounded by approximately 15 m tall trees to the west, the ground dropping away at the other points. The two study sites around the Penryn campus were situated on a rugby pitch (50.166217 N, −5.1192702 W) and in a park in Falmouth (50.157906 N, −5.0783014 W) in the full sun light between the hours of 1100 and 1600 (Figure 1a). The rugby pitch site measures 90 m x 60 m, with approximately 10 m tall trees to the NW, a 5 m high building to the west, a 5 m high bank to the east, and further fields to the south. The release point at this site was at least 40 m away from any of the above features. The park site measured 90 m x 180 m and is situated on a hill. To the west approximately 10 m tall trees are located and houses to the south and north, to the east the ground drops away. Hoverflies were released from the crest of the hill to give them the least obstructed view. At all the locations the migratory insects had a clear, unobstructed view of the sky.

### (d) Assessing reproductive state of migratory hoverflies

Hoverflies were examined in the field by eye to estimate reproductive state based on the fullness of the abdomen. In addition, the reproductive state of two *S. vitripennis* females was examined under a dissection microscope.

### (e) Meteorological recordings

Meteorological conditions were recorded on location at the same time of the experiments. The on-location measurements were cloud cover (OKTA scale), sun visibility, windspeed (Beaufort scale) and wind direction.

### (f) Visualisation of winds

To estimate the potential flight origin, we used Ventusky (www.ventusky.com) to visualise winds at 500m altitude above sea level on the days leading up to the largest influxes.

### (g) Statistical analysis

Statistical analysis and graphing were carried out in R v. 3.5 (13) using R Studio 1.3 and the circular package (14). A Rayleigh test was used to analyse the vanishing bearings of all migratory hoverflies and a Mardia-Watson-Wheeler test was used to look for differences in the distributions of wind and insect vanishing bearings, and between the vanishing bearings of *S. vitripennis* females and all other hoverfly migrants. All data are provided as supplementary material in file S1.

## 3. Results

Released when the sun was visible, hoverflies headed almost due north (θ = 2.3° Rayleigh test: r = 0.59, *p* = >0.0005, n=41, 95% CI 343.5° - 21.1°, Figure 1a, c) after an average orientation period of 2.2 seconds (n = 30). Windspeed during recordings when the sun was visible ranged from 0 to 6 m/s (mean = 1.5 m/s). Under zero-wind conditions, hoverflies headed in a north-northeasterly direction (θ = 18.31° Rayleigh test: r = 0.65, *p* = <0.0005, n = 24, 95% CI 1.7° - 43.6°, Figure 1d). There was no significant difference between the distribution of these vanishing bearings (Mardia-Watson-Wheeler test: W = 35.671, *p* = 0.3). When wind and sun were present simultaneously, the wind direction was either from the west (270°, n=6), the south-southeast (180° to 134°, n=11) or the east (90°, n=1). Under these mixed windy conditions hoverflies headed on average to the north-northwest (θ = 340°, Rayleigh test: r = 0.5981, *p* = <0.0005, n = 18, Figure 1e). Due to low replicates under westerly and easterly wind conditions we were unable to test for an effect of wind direction on hoverfly vanishing bearings during sunny conditions. However, there was no significant difference between the distribution of south-southeasterly wind headings and vanishing bearings under these conditions (Mardia-Watson-Wheeler test: W = 23, *p* = 0.28). Released when the sun was obscured by a thick sea mist, and with a southwesterly wind (238°) blowing at 1.9 m/s, hoverflies headed in a northeasterly direction (θ = 35°, Rayleigh test: r = 0.67, *p* = <0.0005, n = 25, 95% CI 14.7° - 55.2°, Figure 1f). There was no significant difference between the distribution of wind headings and vanishing bearings under these conditions (Mardia-Watson-Wheeler test: W = 47.732, *p* = 0.3236).

Mixed sexes and species of migratory hoverflies were analysed in this study. All females examined showed abdomens without significant egg development and dissections of two female *S. vitripennis* confirmed their pre-reproductive state. Of the 66 hoverflies analysed, 45 were *S. vitripennis* (22 females, 23 males), two were *S. ribesii* (two females), 13 were *E. balteatus* (5 females, 7 males), four were *Scaeva pyrastri* (two females, two males), and two were *Eristalis tenax* (two males). A Mardia-Watson-Wheeler test indicated a lack of significant directional bias by species and sex when comparing the distribution of *S. vitripennis* females with the remainder of the individuals (Mardia Watson Wheeler test: W = 1.4892, *p* = 0.47). To estimate the potential origin of the migratory hoverflies, we visualised the wind conditions at 500 m above sea level at 10:00, 13:00, and 16:00 on the day proceeding the largest influx. These wind conditions suggest an origin for these individuals in France (Figure 2).

**Figure 2.**
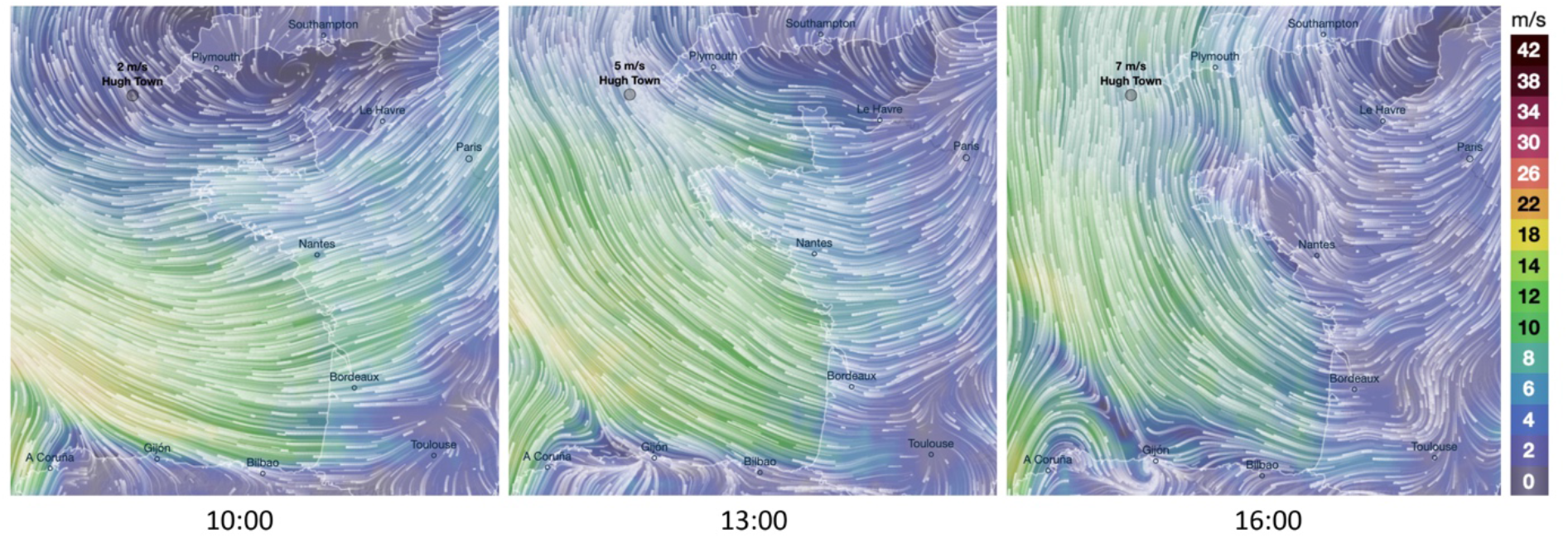
Visualisation of wind directions showing the wind headings on the 16th of June at 10:00, 13:00, and 16:00 at 500m altitude above sea level. Colour gradients signify wind speeds in m/s. Hugh Town on the Island of St Mary’s in the Isles of Scilly is labelled.

## 4. Discussion

The utilisation of favourable winds has been investigated using radar studies in migratory hoverflies (8). These studies suggest a higher selectivity of favourable wind directions during spring mass migrations than in autumn and that, unlike in the autumn, springtime hoverflies do not attempt to correct for wind drift and instead orientate themselves with the downwind heading to increase displacement speed (8). We show here that during a springtime arrival of insect migrants onto the Isles of Scilly and mainland Cornwall, migratory hoverflies orientate and fly in a northerly direction on both sunny and overcast days. Importantly, on sunny days with no wind, hoverflies also headed in a north-northeasterly direction, demonstrating that they are capable of selective orientation during the springtime, rather than simply following the wind. In addition, while the hoverflies were found to fly in a direction not significantly different to the favourable winds, upon release the hoverflies appeared to spend time orientating themselves before leaving, suggesting a period of actively choosing a preferred direction using an internal compass system.

Flight simulator experiments in autumn migrant hoverflies show that they fail to orientate south when the sun is obscured indicating that other available cues are not utilised, at least in this experimental set up (11). Surprisingly, here we find that spring hoverflies also orientate to the north when the sun is obscured, leaving a question as to the cue being used for orientation under these conditions. One explanation may be that hoverflies are simply orientating in a downwind direction. In support of this we note that the more easterly vanishing bearings of the hoverflies are consistent with the southwesterly winds during this experiment. Future experiments under different wind conditions or manipulating other potential direction givers are needed to distinguish between these possibilities.

Our analysis of winds suggests that migrants arriving on the Isles of Scilly began their journey in western France, representing a minimum sea crossing of nearly 200 km (Figure 2). Radar studies indicate spring hoverflies orientate down-wind to increase displacement speed and show significantly faster speeds than autumn migrants with an average of 11.2 m/s (8). We note very similar speeds from our wind analysis that would suggest this distance could be travelled in 5 hours, underlying the importance of warm southerly winds for spring recolonisation of northerly latitudes.

The insect migration assemblage is highly diverse, and the findings presented here on orientation behaviour and migratory routes undoubtedly extend to other members. Many of these migratory insects play important ecological roles (5,6,15), therefore understanding routes and orientation mechanisms used in the spring provides valuable information to understand and predict migration and to benefit from and protect the large-scale movements of these insects.

## Supporting information

FileS1_vanishing bearings data

## Acknowledgments and funding statement

We would like to thank Jason Chapman for his comments on the manuscript. This work was supported through grants to K.R.W from the Royal Society University Research Fellowship scheme (grant no. URF\R\211003). T.D and W.L.S.H were supported by awards to K.R.W from Royal Society Fellows Enhancement Awards (RGF\EA\180083 and RF\ERE\210114). R.M was supported through the NERC GW4+ Doctoral Training Partnership.

## References

1. Hu G, Lim KS, Horvitz N, Clark SJ, Reynolds DR, Sapir N, et al. Mass seasonal bioflows of high-flying insect migrants. Vol. 354, Science. 2016. p. 1584–7.

2. Florio J, Verú LM, Dao A, Yaro AS, Diallo M, Sanogo ZL, et al. Diversity, dynamics, direction, and magnitude of high-altitude migrating insects in the Sahel. Sci Rep. 2020;10(1):1–14.

3. Dingle H. Migration: the biology of life on the move. Oxford University Press, USA; 2014.

4. Chapman JW, Reynolds DR, Wilson K. Long-range seasonal migration in insects: mechanisms, evolutionary drivers and ecological consequences. Ecol Lett. 2015;18(3):287–302.

5. Doyle T, Hawkes WLS, Massy R, Powney GD, Menz MHM, Wotton KR. Pollination by hoverflies in the Anthropocene: Pollination by Hoverflies. Vol. 287, Proceedings of the Royal Society B: Biological Sciences. 2020.

6. Satterfield DA, Sillett TS, Chapman JW, Altizer S, Marra PP. Seasonal insect migrations: massive, influential, and overlooked. Frontiers in Ecology and the Environment. 2020.

7. Wotton KR, Gao B, Menz MHM, Morris RKA, Ball SG, Lim KS, et al. Mass Seasonal Migrations of Hoverflies Provide Extensive Pollination and Crop Protection Services. Vol. 29, Current Biology. 2019. p. 2167-2173.e5.

8. Gao B, Wotton KR, Hawkes WLS, Menz MHM, Reynolds DR, Zhai BP, et al. Adaptive strategies of high-flying migratory hoverflies in response to wind currents. Vol. 287, Proceedings of the Royal Society B: Biological Sciences. 2020. p. 20200406.

9. Oliveira EG, Srygley RB, Dudley R. Do neotropical migrant butterflies navigate using a solar compass? Journal of Experimental Biology. 1998;201(24):3317–31.

10. Schmidt-Koenig K. Directions of migrating monarch butterflies (Danaus plexippus; Danaidae; Lepidoptera) in some parts of the eastern United States. Behavioural processes. 1979;4(1):73–8.

11. Massy R, Hawkes WLS, Doyle T, Troscianko J, Menz MHM, Roberts NW, et al. Hoverflies use a time-compensated sun compass to orientate during autumn migration. Proceedings of the Royal Society B. 2021;288(1959):20211805.

12. Mouritsen H, Frost BJ. Virtual migration in tethered flying monarch butterflies reveals their orientation mechanisms. Vol. 99, Proceedings of the National Academy of Sciences of the United States of America. 2002. p. 10162–6.

13. R Core Team. R: A language and environment for statistical computing.. Vienna, Austria.: R Foundation for Statistical Computing; 2020.

14. Agostinelli C, Lund U. R package ‘circular’: circular statistics (version 0.4-93). URL https://r-forger-projectorg/projects/circular. 2017;

15. Hawkes WLS, Walliker E, Boya G, Forster O, Lacey K, Doyle T, et al. Huge spring migrations of insects from the Middle East to Europe: quantifying the migratory assemblage and ecosystem services. Ecography. 2022;

